# The accumulation of orphan ribosomal proteins is a hallmark of ALS

**DOI:** 10.64898/2026.05.18.725994

**Authors:** Anabel Sáez-Mas, Guillermo de la Vega-Barranco, Amin El-Manchoud, Iván Ventoso, Sara Rodrigo, Vanesa Lafarga, Oscar Fernández-Capetillo

## Abstract

Amyotrophic lateral sclerosis (ALS) is a progressive neurodegenerative disease of poor prognosis, for which age is the strongest risk factor. Despite significant progress in the discovery of ALS-associated mutations, no model explains how such a diversity of mutations converges in a common pathology. In addition, most ALS cases are sporadic and lack known genetic drivers. We recently reported that arginine-rich peptides arising from the *C9ORF72* mutation trigger a widespread accumulation of orphan ribosomal proteins (oRP). Here, we show that oRP accumulation is also observed upon expression of other RNA-related ALS mutations, such as hnRNPA2^D290V^ and TDP-43^A315T^, as well as upon exposure to the ALS-related neurotoxin β-N-methylamino-L-alanine (BMAA). Furthermore, the transcriptional signature of patients with sporadic ALS resembles that of Diamond-Blackfan anemia (DBA), a known ribosomopathy. Supporting the usefulness of our in vitro data, a transcriptional signature defined from these models provides diagnostic and prognostic value in ALS patients. We propose that the accumulation of oRPs due to dysfunctional ribosome biogenesis is a molecular hallmark of ALS that can contribute to the progressive loss of motor neurons in the disease.

## Introduction

Amyotrophic lateral sclerosis (ALS) is a fatal neurodegenerative disorder for which there are few therapeutic options available^1,2^. As for most neurodegenerative conditions, age remains the strongest risk factor^3^. While the majority of ALS cases are sporadic and thus of unknown etiology, significant progress has been made in the discovery of ALS-associated mutations. Mutations in SOD1 were first to be found and drove significant attention to the role of oxidative stress in the disease^4,5^. However, targeting oxidative damage has shown limited benefits in ALS patients^6^. Subsequent to SOD1, ALS-associated mutations were often found in genes encoding RNA-binding proteins, such as TIA1, TAF15, ATXN2, SETX, FUS, hnRNPA2 and TAR DNA-binding Protein (TDP-43)^7,8^. Furthermore, loss of nuclear localization and/or cytosolic aggregates of TDP-43 arguably constitute the most widespread molecular hallmark of ALS and the related frontotemporal dementia (FTD)^9,10^. This phenomenon, which has now been documented in additional neurodegenerative conditions, has been associated with perturbations that impair TDP-43 interactions with RNA that occur inside the nucleus^11–14^.

Despite the growing list of mutations associated with ALS, a unifying mechanism for this disease remains elusive. Currently, no model can explain how such a wide diversity of mutations converges on a common cellular pathology. The high prevalence of sporadic cases further complicates this landscape, highlighting the need to identify the underlying cellular dysfunction that drives motor neuron degeneration. Nevertheless, the high frequency of mutations affecting RNA-binding proteins has long suggested a prevailing role for dysfunctional RNA metabolism in the origins of ALS^15,16^.

This RNA-centric view of ALS is further supported by the most frequently mutated gene, *C9ORF72.* Although precise numbers fluctuate in different populations, *C9ORF72* mutations account for roughly one-third of familial ALS cases and 4-7% of sporadic cases^17–19^. The mutation involves the expansion of a GGGGCC repeat within the first intron of the gene, which reduces its expression, forms toxic RNA foci, and undergoes repeat-associated non-AUG (RAN) translation to produce dipeptide repeats (DPRs), including the highly toxic arginine-rich (GR)n and (PR)n species^17,19–24^. Interestingly, arginine-rich DPRs perturb nucleolar function, RNA biogenesis, and the dynamics of RNA granules, further indicating that alterations of RNA metabolism play a significant role in C9ORF72-associated neurodegeneration^22,25,26^.

Initial work on *C9ORF72*-associated arginine-rich peptides revealed that these accumulated at nucleoli and perturbed rRNA maturation^22^. Subsequent studies from our laboratory demonstrated that, due to the high affinity of arginines for RNA, these peptides decorate all cellular RNAs and trigger a generalized displacement of RNA-binding proteins (RBPs) from RNA^27^. Furthermore, their impact on rRNA biogenesis leads to a generalized accumulation of orphan ribosomal proteins (oRPs)^28^. Because oRPs are highly basic and disordered when not bound to rRNA, they are prone to rapid aggregation, triggering a proteotoxic response that compromises viability in eukaryotic cells and animal models^29–31^. Building on these discoveries, we here set to address whether dysfunctional rRNA biogenesis and oRP accumulation are a shared hallmark of ALS.

## Results

### ALS mutations impair rRNA processing, leading to mTOR hyperactivation

To investigate the cellular consequences of expressing ALS mutations that are related to RNA biology, we used three U2OS cell lines engineered to enable doxycycline (dox)-dependent expression of TDP43^A315T^, hnRNPA2^D290V^ or (PR)_97_ peptides^32^. First, we evaluated whether the expression of these mutations affected rRNA processing. To do so, we performed fluorescent in situ hybridization (FISH) using rRNA-directed probes and quantified the signal by High-Throughput Microscopy (HTM) (**Fig. 1A**). Consistent with previous work using recombinant (PR)n peptides^22^, dox treatment impaired rRNA maturation in U2OS^PR97^ cells as evidenced by the accumulation of 47S pre-rRNA at nucleoli, which was also observed upon expression of TDP43^A315T^ or hnRNPA2^D290V^ (**Fig. 1B-D**). This nucleolar accumulation or rRNA precursors was not due to increased rRNA expression which, to the contrary, was reduced in all cases (**Fig. S1A**). Of note, a buildup of rRNA precursors can account for the enlarged nucleolar area previously reported in motorneurons from ALS patients^33^, and which we also observe upon expression of TDP43^A315T^, hnRNPA2^D290V^ or (PR)_97_ peptides (**Fig. S1B**). Together with the accumulation of rRNA precursors in the nucleolus, TDP43^A315T^, hnRNPA2^D290V^ and (PR)_97_ expression led to lower cytoplasmic levels of mature 28S (**Fig. 1B-D**). These results indicate that defective rRNA maturation is a common phenotype of cells expressing RNA-related ALS mutations.

**Figure 1.**
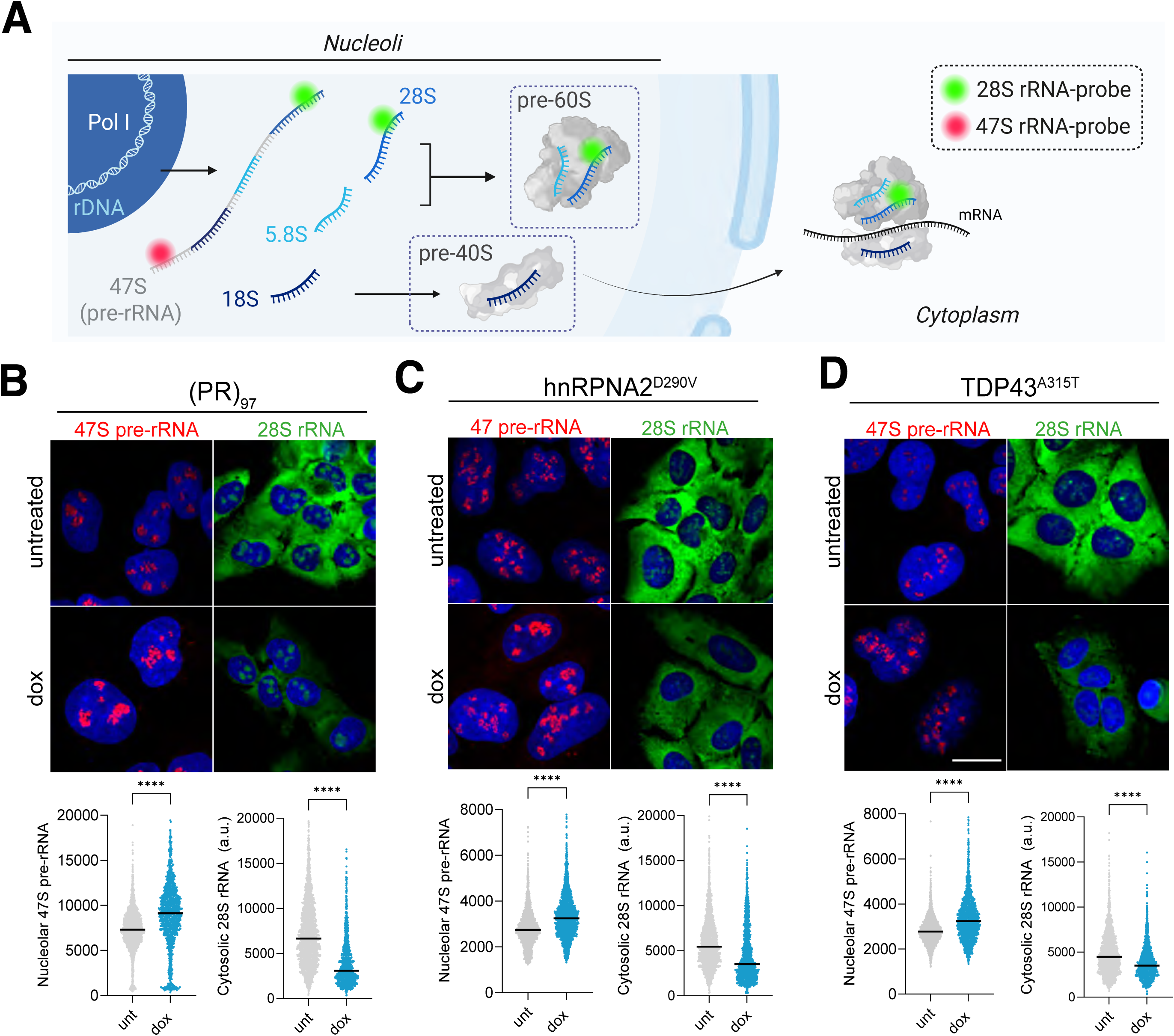
RNA-related ALS mutations impair rRNA processing. (**A**) Schematic illustration of rRNA processing steps and hybridization using 47S pre-rRNA and 28S rRNA probes. (**B-D**) Representative images of 47S (red) and 28S (green) signals from in situ hybridizations in untreated and dox treated (48h) U2OS cells expressing (PR)_97_ (**B**), hnRPNA2^D290V^ (**C**) or TDP43^A315T^ (**D**). DAPI (blue) was used to stain DNA. Scale bar (white) indicates 10 μm. HTM-mediated quantification of the signal intensities is shown in bottom panels. Black lines indicate median values. The graph represents one experiment from 3 biological repeats. \*\*\*\**P* < 0.0001; *t*-test.

Eukaryotic cells respond to impaired rRNA processing by activation of mTOR signaling, in an effort to restore ribosome biogenesis^34^. Accordingly, western blotting (WB) revealed increased phosphorylation of the mTOR target RPS6 in response to TDP43^A315T^, hnRNPA2^D290V^ or (PR)_97_ expression (**Fig. S1C**). Moreover, and as previously shown for (PR)_97_ peptides in vitro and in vivo^28^, mTOR inhibition with rapamycin alleviated the toxicity associated to (PR)_97_, TDP43^A315T^ or hnRNPA2^D290V^ expression (**Fig. S1D**). Collectively, these data indicate that ALS mutations related to RNA biology impair rRNA processing, triggering a compensatory toxic hyperactivation of the mTOR pathway.

### Accumulation of orphan RPs is a common hallmark of ALS mutations

Next, we set to determine the consequences of mTOR activation in cells expressing RNA-related ALS mutations. One of the main outputs of mTOR activity is the expression of ribosomal proteins (RPs), which occurs through activation of the transcription factor c-MYC^35^ or by selectively stimulating the translation of RP mRNAs^36^. Consistently, expression of TDP43^A315T^, hnRNPA2^D290V^ or (PR)_97_ led to a significant increase in cytoplasmic RPS6 levels, as quantified by HTM (**Fig. 2A**). Given that RP accumulation occurs in the context of reduced levels of mature rRNA, we hypothesized that this should lead to oRP accumulation.

**Figure 2.**
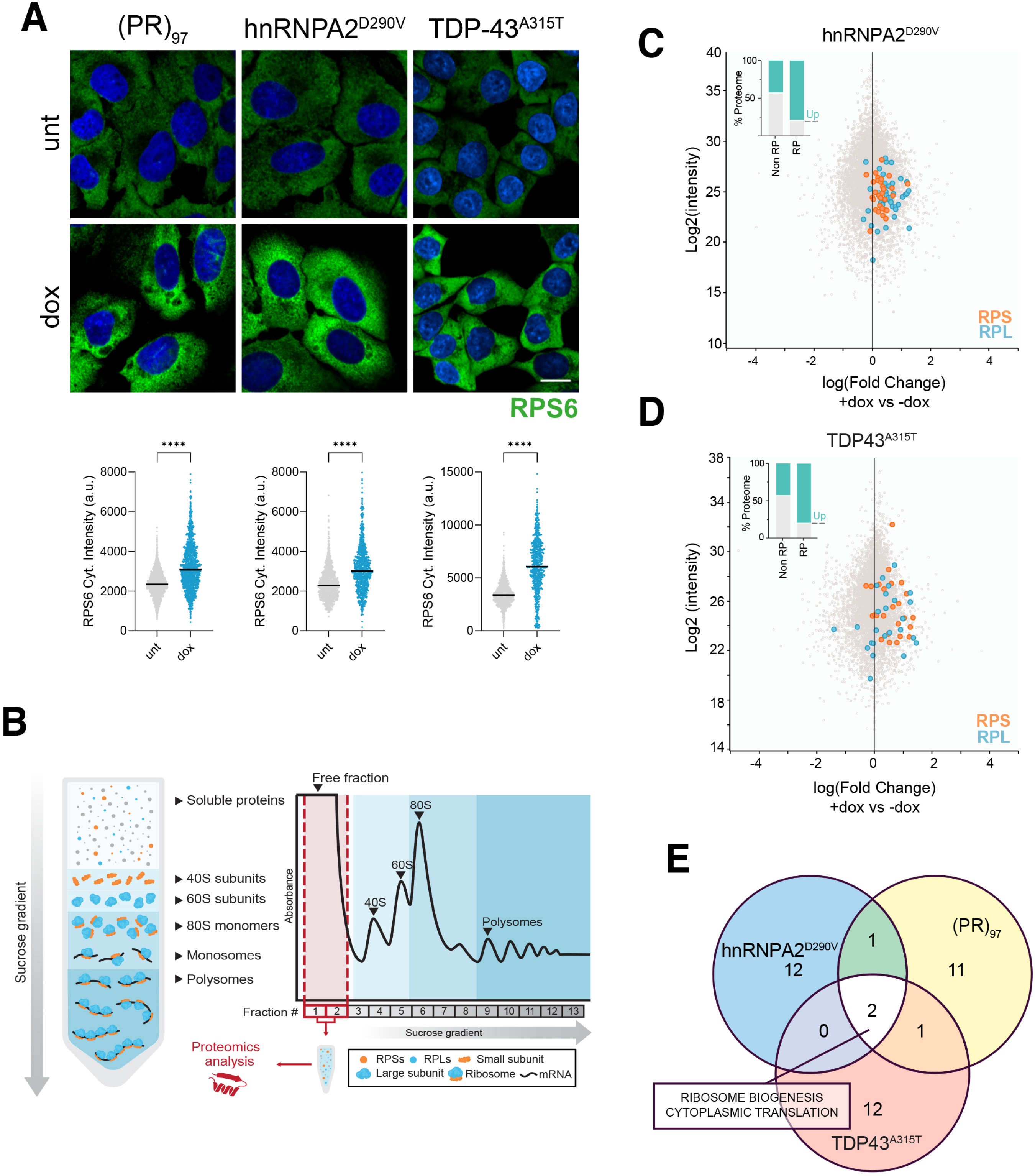
Expression of RNA-related ALS mutations triggers oRP accumulation. (**A**) Representative images of RPS6 immunofluorescence (green) in untreated and dox treated U2OS cells expressing (PR)_97_, hnRPNA2^D290V^ or TDP43^A315T^. DAPI (blue) was used to stain DNA. Scale bar (white) indicates 10 μm. HTM-mediated quantification of the signal intensities is shown in bottom panels. Black lines indicate median values. The graph represents one experiment from 3 biological repeats. Data information: \*\*\*\**P* < 0.0001; *t*-test. **(B)** Schematic illustration of polysome profiling and sub-fractionation used for proteomic analyses. Fractions 1 and 2, corresponding to the “Free fraction” (soluble proteins not forming part of complexes), were pooled and processed for downstream quantitative proteomics. (**C-D**) MA plot representing the Log_2_(intensity) against log_2_(fold-change) of all proteins identified by proteomics of the ribosome-free fraction of dox-treated (48 hrs) U2OS cells expressing hnRPNA2^D290V^ (**C**) or TDP43^A315T^ (**D**), compared to untreated cells. RPs are marked with orange (40S) and blue (60S). Inset: Stacked bar chart summarizing the gross distribution of (RPs) and non-ribosomal proteins (non-RPs). **(E)** Venn Diagram illustrating the overlap of the top 15 significantly upregulated Gene Ontology (GO) Biological Processes (*P* < 0.01, ranked by NES) derived from proteomic data of U2OS cells expressing (PR)_97_, hnRPNA2^D290V^ and TDP43^A315T^. Proteomic data of the soluble fraction of U2OS cells (PR)_97_ expressing were previously reported^28^.

To evaluate oRP abundance, we isolated polysomes using a sucrose gradient fractionation protocol. We then analyzed the free fraction by proteomics, which contains individual proteins that are not forming part of larger complexes (**Fig. 2B**). Consistent with our previous findings in response to (PR)_97_ peptides^28^, expression of TDP43^A315T^ or hnRNPA2^D290V^ led to a widespread accumulation of oRPs (**Fig. 2C, D**). In fact, Gene Set Enrichment Analyses (GSEA) using these proteomic data revealed that “Ribosome Biogenesis” and “Cytoplasmic Translation” were the most significantly upregulated biological pathways across all mutations (**Fig. 2E**). Together, these results suggest that oRP accumulation is a shared hallmark of cells expressing RNA-related ALS mutations.

### Molecular resemblance between ALS mutations and ribosomopathies

Next, we used our proteomic data to interrogate the “DISEASES” signatures from the laboratory of Lars J. Jensen^37^, in order to identify human diseases that have an expression profile that resembles the one observed upon TDP43^A315T^, hnRNPA2^D290V^ or (PR)_97_ expression. Strikingly, the disease showing the highest significant similarity to hnRNPA2^D290V^ or (PR)_97_ expression, and third to TDP43^A315T^, was “Diamond Blackfan Anemia” (DBA), a congenital ribosomopathy^38^ (**Fig. 3A**). Furthermore, DBA and Pure Red Cell Aplasia (PRCA), both related to ribosome dysfunction, were the only two diseases with significant similarity that were shared among the three cellular models (**Fig. 3B**). Noteworthy, oRP accumulation is the defining feature of ribosomopathies, hereditary disorders associated to mutations in ribosomal proteins. We thus hypothesized that our observations upon expression of ALS mutations should also be present in ribosomopathy models.

**Figure 3.**
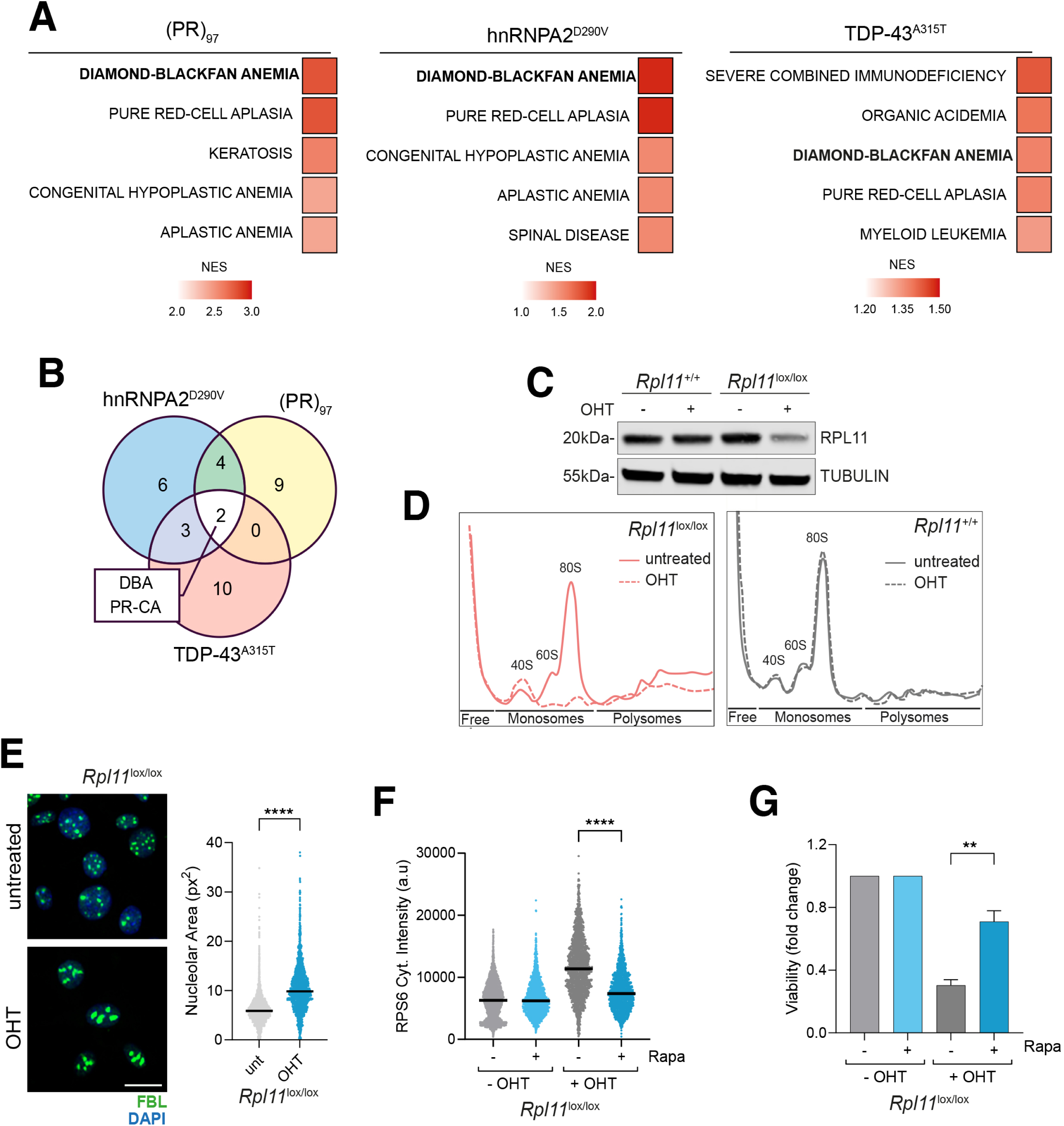
Similarities between ALS and ribosomopathies. (**A**) GSEA analyses using proteomic data of U2OS cells expressing (PR)_97_, hnRPNA2^D290V^ and TDP43^A315T^. Analyses were run against the “DISEASES” database of expression signatures from human diseases developed by the Jensen laboratory^37^. Heatmaps display the top five disease terms ranked by NES. Colour intensities represent NES values. (**B**) Venn Diagram illustrating the overlap of the 15 most significantly associated diseases across the three proteomic datasets (DBA: Diamond Blackfond anemia; PR-CA: Pure red cell aplasia). (**C**) WB of RPL11 levels in *Rpl11*^+/+^ and *Rpl11*^lox/lox^ MEFs expressing Cre-ERT2 after 4-OHT treatment (72h). TUBULIN levels are shown as a loading control. (**D**) Representative polysome profiles of *Rpl11*^+/+^ (red) and *Rpl11*^lox/lox^ (black) MEFs untreated (solid line) or treated with 4-OHT (dashed line) for 72h. (**E**) Representative immunofluorescence of the nucleolar factor FBL (green) in *Rpl11*^lox/lox^ MEF upon 4-OHT treatment (48h). DAPI (blue) was used to stain DNA. Scale bar (white) indicates 10 μm. HTM-mediated quantification of the signal intensities is shown in the right panel. (**F**) HTM-mediated quantification of cytoplasmic RPS6 levels in *Rpl11*^lox/lox^ MEF upon 4-OHT treatment (48h), in the presence or absence of rapamycin (Rapa, 50 nM). (**G**) Cell viability, measured by HTM-mediated quantification of nuclei in *Rpl11*^lox/lox^ MEF upon 4-OHT treatment (48h), in the presence or absence of rapamycin (Rapa, 50 nM). \*\*\*\**P* < 0.0001; \*\**P* < 0.01; *t*-test.

To address this, we used a previously developed mouse model of ribosomopathy based on the conditional depletion of RPL11 (*Rpl11*^lox/lox^)^39^. As expected, RPL11 depletion led to generalized loss of the 60S and 80S ribosomal complexes in mouse embryonic fibroblasts (MEF) from these mice (**Fig. 3C, D**). Significantly, this loss of mature ribosomes was associated with an increase in the nucleolar area (**Fig. 3E**) and in cytoplasmic RPS6 levels (**Fig. 3E, F**). We reasoned that these findings could once again be the result of a futile compensatory activation of mTOR signaling, that is aiming to restore ribosome biogenesis. Accordingly, treatment with rapamycin abrogated the increase in RPS6 levels found in RPL11-deficient MEF, and improved their overall viability (**Fig. 3F, G**). In agreement with our findings, rapamycin was previously shown to be beneficial in a DBA-model based on hematopoietic progenitors^40^, and in a Drosophila model of ribosomopathy^30^. Together, these results indicate that ALS mutations drive a toxic accumulation of oRPs, which resembles the one found in human ribosomopathies.

### Transcriptomic convergence across ALS and ribosomopathy models

As mentioned, mTOR activation leads to increased RP transcription. We thus hypothesized that the transcriptional landscape of cells suffering from oRP accumulation should include a generalized increase in RP mRNA levels. To test this, we first performed RNAseq in RPL11-deficient MEF. As predicted, and besides the expected loss of *Rpl11* expression, there was a widespread upregulation of RP mRNA expression in the mutant cells (**Fig. 4A**). Furthermore, the most significantly upregulated pathways found in RPL11-deficient MEF, as detected by GSEA, were primarily related to ribosome biogenesis and rRNA processing (**Fig. 4B**). Consistent with MEF data, a meta-analysis of RNAseq from human DBA cells revealed a generalized upregulation of RP transcription (**Fig. 4C**).

**Figure 4.**
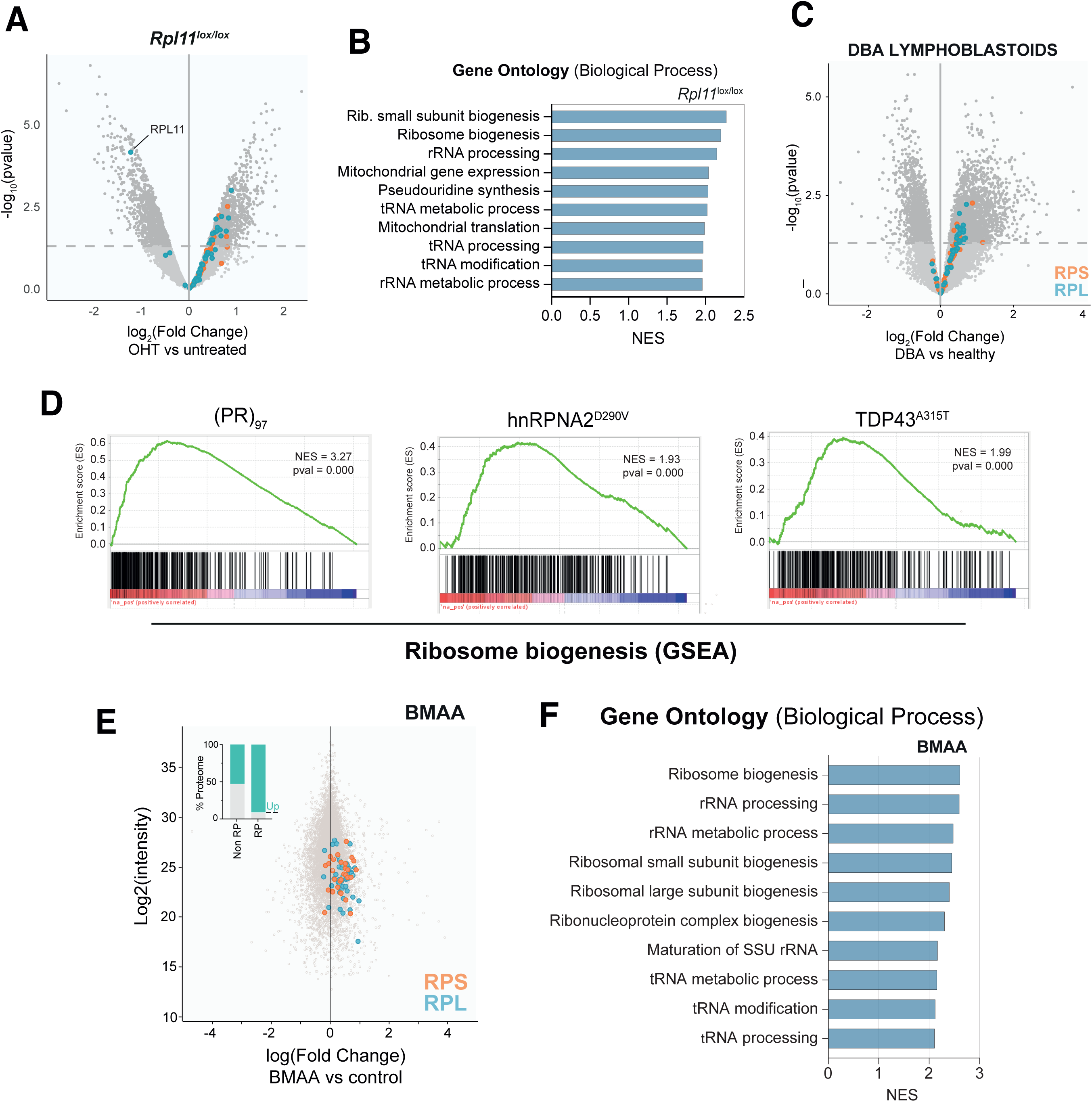
Increased rRNA processing and ribosome biogenesis in ALS and DBA cell models. (**A**) Volcano plot displaying differentially expressed genes obtained from RNAseq data of *Rpl11*^lox/lox^ MEF treated with 4-OHT for 48h compared to untreated controls. Log2(fold change) is plotted against −;log_10_(*P* value). The downregulation of *Rpl11* is highlighted, along with the levels of RPs (RPL: blue; RPS: orange). (**B**) Top significantly upregulated GO Biological Processes (*P* < 0.01) ranked by NES from RNAseq data of *Rpl11*^lox/lox^ MEF shown in (**A**). (**C**) Volcano plot displaying differentially expressed genes obtained from RNAseq data of lymphoblastoid cells from DBA patients compared to healthy controls. Log2(fold change) is plotted against −;log_10_(*P* value). RP expression is highlighted (RPL: blue; RPS: orange). (**D**) GSEA enrichment plots of the GO Biological Process ‘Ribosome Biogenesis’ from RNAseq data obtained from U2OS cells expressing (PR)_97_, hnRPNA2^D290V^ or TDP43^A315T^. NES and *P* values are indicated. (**E**) MA plot displaying proteomic data of the soluble fraction from U2OS cells treated with 20 mM L-BMAA for 24 h. Log2(intensity) is plotted against Log2(fold change) relative to untreated controls. RP levels are highlighted (RPL: blue; RPS: orange). Inset: Stacked bar chart summarizing the distribution of RP and non-RP abundances. (**F**) Top significantly upregulated GO Biological Processes (*P* < 0.01) ranked by NES from RNAseq data of U2OS cells treated with 20 mM L-BMAA for 24 h.

Next, we explored if similar observations were made in the context of ALS mutations. To do so, we performed RNAseq in U2OS cells expressing (PR)_97_, TDP43^A315T^ or hnRNPA2^D290V^. Consistent with our findings in ribosomopathy, GSEA analyses revealed that all models presented a significant upregulation in “ribosome biogenesis” (**Fig. 4D**). Given that most cases of ALS are sporadic with no associated mutation, we also wanted to investigate the effect of environmental factors. In this regard, one of the best examples is that of β-N-methylamino-L-alanine (BMAA), a non-proteinogenic aminoacid produced primarily by cyanobacteria. Chronic dietary exposure to BMAA triggers a complex neurodegenerative pathology with symptoms related to ALS and parkinsonism-dementia complex (ALS-PDC)^41,42^. Remarkably, BMAA exposure led to a widespread accumulation of oRPs (**Fig. 4E**), as detected by proteomics, together with the transcriptional upregulation of pathways related to ribosome biogenesis and rRNA processing, as detected by RNAseq (**Fig. 4F**). Hence, ALS-associated mutations and environmental factors trigger a widespread accumulation of oRPs together with a transcriptional upregulation of ribosome biogenesis.

### Upregulation of ribosome biogenesis in ALS patients

Next, to evaluate if our findings from *in vitro* models were also observed in ALS patients, we interrogated RNAseq data available at the New York Genome Centeŕs ALS Consortium that is based on 189 ALS patients (149 sporadic) and 46 controls (**Fig. S2A**). Early analyses of this dataset focused on inflammatory aspects of the disease and revealed a significant increase in activated microglia and astrocytes in ALS biopsies^43^. Consistently, Gene Ontology analyses using the “Biological Process” category, shows an enrichment of inflammatory pathways in the spinal cord of ALS patients (**Fig. 5A, B**). However, when Gene Ontology searches were performed using the term “Cellular Component”, the most significantly upregulated pathways were primarily related to ribosome biology (**Fig. 5C**). Moreover, GSEA enrichment plots showed a significant upregulation of “ribosome biogenesis” and “rRNA processing” in ALS spinal cords (**Fig. 5D**). Importantly, the transcriptome of ALS patients presented a significant upregulation of RP expression in affected areas such as the spinal cord, but which was absent in non-affected tissues such as the cerebellum (**Fig. 5E, F**).

**Figure 5.**
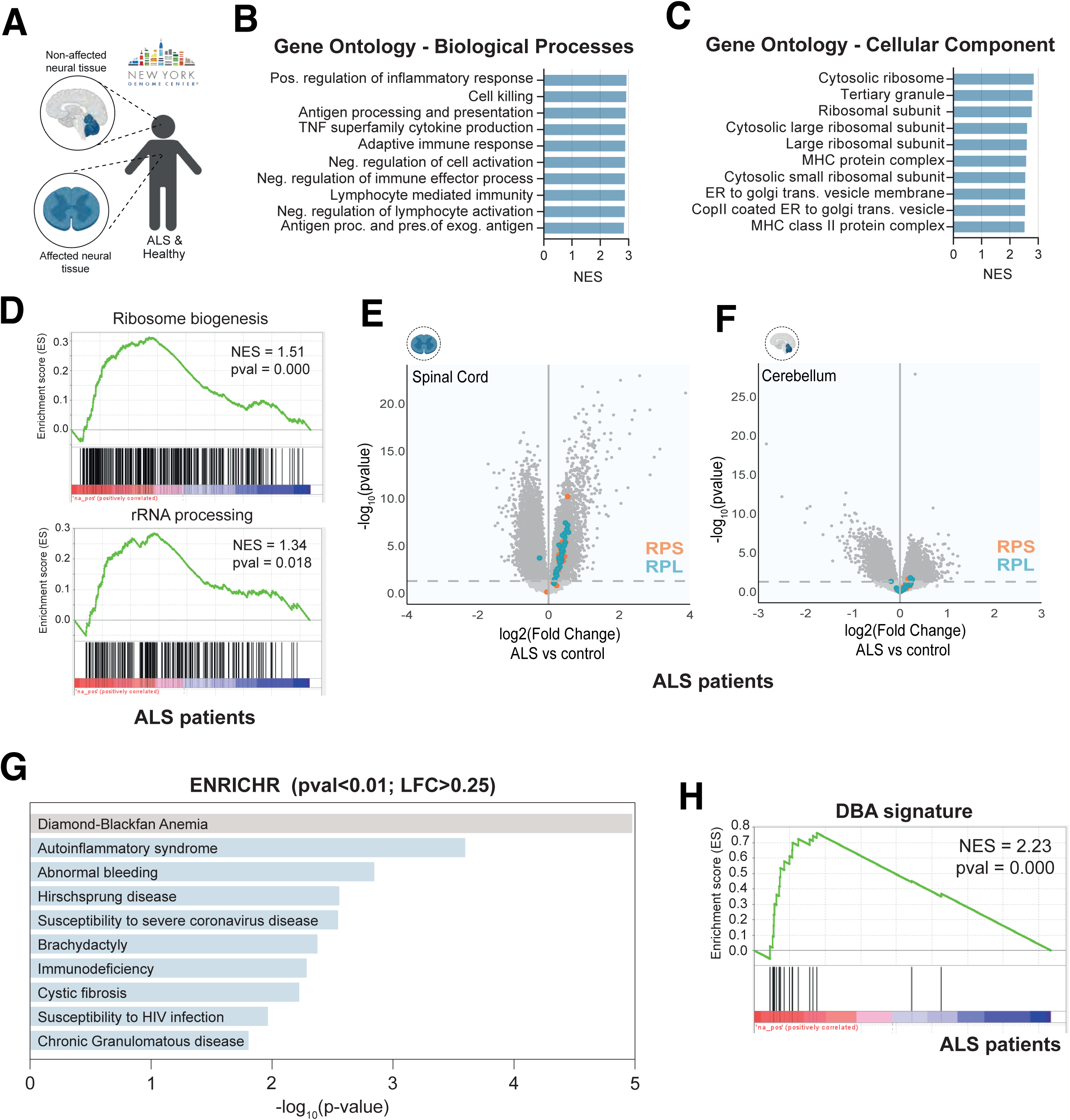
Upregulation of rRNA processing and ribosome biogenesis in ALS patients. (**A**) Schematic representation of samples included in the study from the New York Genome Center. The cohort includes RNAseq data from 189 ALS patients (149 sporadic) and 46 controls. (**B-C**) Top 10 significantly upregulated GO terms for “Biological Processes” (**B**) and “Cellular Components” (**C**), from genes significantly overexpressed in ALS patients, ranked by NES. (**D**) GSEA enrichment plots for “Ribosome Biogenesis” and “rRNA processing” using RNAseq data from ALS patients (vs controls) defined in (**A**). (**E-F**) Volcano plots displaying differentially expressed genes using RNAseq data of ALS patients from affected tissue (cervical spinal cord (**E**)) and unaffected tissue (cerebellum (**F**)), compared to healthy controls. Log_2_(fold change) is plotted against −;log_10_(*P* value). RP expression is highlighted (RPL: blue; RPS: orange). (**G**) Over-representation analysis (ORA) using the ENRICHR platform of pathway analyses (*P* < 0.01, Log_2_FC > 0.25). The bar graph displays the top 10 clinical conditions that have a transcriptional signature that is most similar to the one from the cervical spinal cord dataset of ALS patients. (**H**) GSEA enrichment plot for the “DBA signature” using RNAseq data from ALS patients (vs controls).

Next, and using the data from the NYGC ALS Consortium, we selected the genes that were significantly overexpressed on the spinal cord of ALS patients. We then used this list to evaluate its similarity to the transcriptional profile of other human diseases, by comparison against curated disease-associated gene lists available at ENRICHR^44^. Strikingly, and consistent with our analyses using proteomic data, these studies revealed that the human disease that most significantly resembles ALS at the transcriptional level was, once again, DBA (**Fig. 5G, H**). Collectively, these findings reinforce that dysfunctional ribosome biogenesis is a hallmark of ALS, including in sporadic cases of the disease.

### A transcriptional signature of ALS defined from in vitro expression models

Our previous results indicated that the molecular alterations observed in our cellular models of TDP43^A315T^, hnRNPA2^D290V^ or (PR)_97_ expression, were reminiscent of changes that are present in ALS patients. On this basis, we hypothesized that a transcriptional signature defined from genes showing altered expression in the three models could have predictive value in the actual disease. A selection of overlapping DEGs led to an initial gene signature of 433 genes, which already demonstrated predictive value across most tissues from the NYGC ALS Consortium (**Fig. S2B**). Next, we refined the signature using machine learning to train on RNAseq data from cervical spinal cord tissue of ALS patients, resulting in a final signature of 61 genes (ALS^sig^; **Fig. 6A, B; Table S1**). As expected, when applied to transcriptomic data from cervical spinal cord, ALS^sig^ separated ALS patients from controls with a p-value of p-value of 9.0 × 10^-13^ (**Fig. 6C**). Importantly, the signature also showed strong predictive value in lumbar spinal cord (**Fig. 6C**) and in additional tissues from the NYGC ALS dataset (**Fig. S2B, C**). Interestingly, the signature also separated ALS patients from controls in RNAseq data from blood samples, albeit with reduced significance (**Fig. 6D**), suggesting that the molecular pathology that drives motor neuron loss is also present in unaffected tissues, yet with reduced severity. The effectiveness of our ALS^sig^ was validated in a smaller cohort of spinal cord RNAseq data (8 ALS patients and 4 age-matched controls) (**Fig. 6E**). Finally, we explored if our ALS^sig^ could also have prognostic value. In fact, classifying the patients from the NYGC ALS Consortium dataset on the basis of ALS^sig^, significantly segregated patients with different severity of the disease (defined on the basis of life expectancy from the onset of the disease) (**Fig. 6F**). Of note, the ALS^sig^ defined in this study substantially outperformed than the ones existent in the KEGG database or in the “DISEASES” dataset from the Jensen laboratory (**Fig. 6G, Fig. S2D**). Besides the usefulness of ALS^sig^ as a novel signature of ALS, these analyses support the value of in vitro mutation expression models as valid platforms for unraveling molecular alterations associated to ALS.

**Figure 6.**
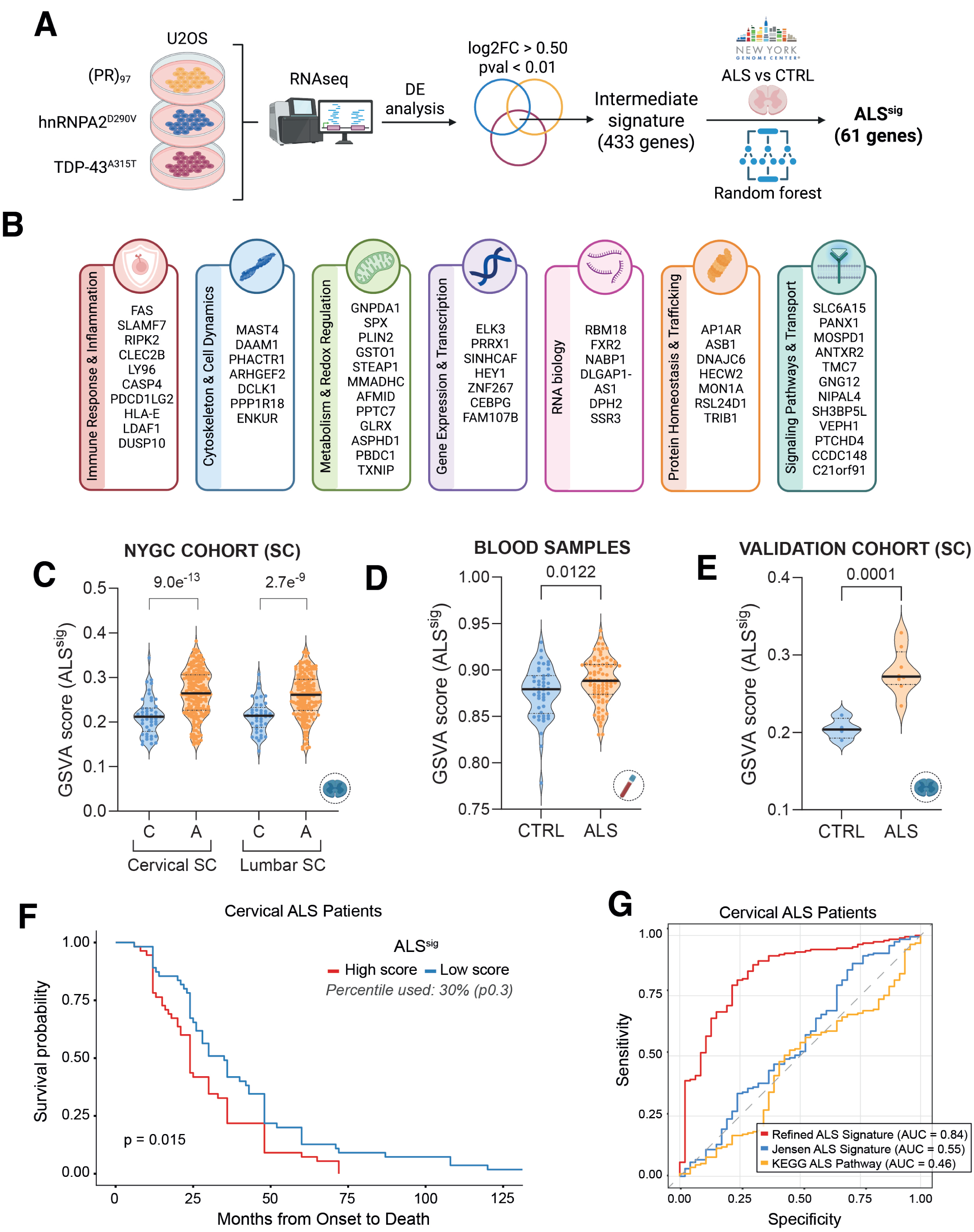
An ALS transcriptional signature defined from mutation expression models. (**A**) Schematic representation of the computational pipeline used to define the transcriptional signature of ALS (ALS^sig^). An initial signature of 433 genes was first defined using common differentially expressed genes (DEGs) from RNAseq data of U2OS cells expressing (PR)_97_, hnRPNA2^D290V^ or TDP43^A315T^. This was followed by dimensionality reduction using Random Forest on the NYGC spinal cord patient dataset to refine into a final 61-gene signature (**Table S1**). (**B**) Functional classification of the ALS^sig^ genes grouped by their primary cellular functions. (**C–E**) Gene Set Variation Analysis (GSVA) scores for the ALS^sig^ across different cohorts: spinal cord samples from the NYGC (**C**; including Cervical training set and Lumbar validation set), peripheral blood samples from ALS patients (**D**), and an independent spinal cord validation cohort (**E**). (**F**) Kaplan-Meier survival analysis of ALS patients stratified by their ALS^sig^ score (high score, red; low score, blue) based on the 30th percentile. (**G**) Receiver Operating Characteristic (ROC) curve comparing the diagnostic performance (AUC) of the ALS^sig^ against established ALS-related signatures from Jensen and KEGG databases using cervical spinal cord samples from the NYGC consortium.

## Discussion

The aggregation of misfolded proteins is a hallmark of most neurodegenerative disorders^45^, and these aggregates frequently involve RBPs. This is also the case in ALS, where mutations in RBPs are the most frequent class, together with mutations that affect proteostasis and macro-autophagy which might impair the clearance of RBP aggregates^46^. These findings suggest that dysfunctional RNA metabolism might be the molecular determinant of the cellular toxicity that drives motorneuron loss in ALS patients. In this regard, substantial evidence supports alterations in various aspects of RNA metabolism such as mRNA transcription, splicing or export in ALS, and therapeutic strategies are being explored to correct these problems^47^. The work presented brings a new perspective to the field, by identifying oRP accumulation as another consequence of RNA-related ALS mutations.

Several reasons suggest that oRP accumulation might play an important role in the toxic processes that promote neurodegeneration in ALS. First, their abundance. In eukaryotic cells the vast majority of RNA, up to 85%, is rRNA and RPs often exceed 40% of the total cellular proteome by mass^48^. Second, and as mentioned above, the basic nature of RPs makes them aggregation prone when not forming part of ribosomes, and these aggregates have been shown to drive cellular toxicity in vitro and in animal models^29–31^. An important consideration to be made is that the detection of oRP aggregates is technically challenging, particularly in clinical samples. Polysome fractionation approaches demand a very high quality of RNA, which is difficult to preserve in autopsies. Nevertheless, since dysfunctional ribosome biogenesis triggers a compensatory transcriptional response, this has enabled us to evaluate these phenomena in ALS samples. Importantly, the fact that a transcriptional signature defined from non-neuronal cellular models has diagnostic and prognostic value in ALS underscores that the fundamental processes that drive the loss of motorneurons in this disease, are shared across cell types.

Another interesting aspect that emerges from our work relates to the relationship between aging and neurodegeneration. Interestingly, a recent study reported that aged neurons present a substantial depletion of RBPs^49^. In addition, rRNA biogenesis has been shown to decline during aging, particularly in post-mitotic cell types such as neurons^50,51^ and RP aggregates have been shown to accumulate in the aging brain^52^. One possibility that emerges from these findings is that, by perturbing rRNA processing and ribosome biogenesis, ALS mutations or environmental factors might be triggering toxic cellular responses that normally occur later in life. In this scenario, ALS could be understood as an accelerated aging of the motorneurons. Consistent with this idea, recent studies are highlighting that human organs age at different rates, and that diseased organs often present the molecular hallmarks of older ones ^53–56^.

To end, our study also suggests that strategies oriented to reduce the abundance of oRPs might be of therapeutic interest in ALS. To do so, one approach could be to slightly reduce ribosome biogenesis rates by targeting key mediators such as mTOR or MYC. Along these lines, our previous data with (PR)_97_ peptides^28^, and the data presented in this study, show that mTOR inhibition can alleviate the cellular toxicity driven by RNA-related ALS mutations. Of note, while a significant reduction in ribosome biogenesis might be deleterious in the long term, milder approaches might yield beneficial effects. Consistent with this notion, a seminal study from the laboratory of Adam Antebi revealed that small nucleoli are associated with longevity, and that dampening nucleolar activity by a partial depletion of FBL extends lifespan in *C. elegans*^57^. In any case, an alternative strategy to limiting ribosome biogenesis could be to promote the clearance of RP aggregates. In strong support of this idea, we have recently reported that stimulating ribophagy, the autophagic clearance of dysfunctional ribosomes, rescues the toxicity of (PR)n peptides in cells (including neurons) and in a *Drosophila* model of *C9ORF72* ALS^58^. In summary, the work presented here supports that dysfunctional ribosome biogenesis and oRP accumulation are convergent molecular hallmarks of ALS and suggests that strategies oriented to reduce oRP aggregates might be an innovative therapeutic approach for the treatment of this fatal disease.

## Materials and Methods

### Cell Culture

Cell viability was quantified by HTM as total nuclei counts per well relative to controls. Cells were seeded in μCLEAR-bottom 96-well plates (Greiner Bio-One) at 3,000–6,000 cells/well (as specified per experiment). Fixation was performed with 4% paraformaldehyde (PFA) in PBS for 10 min at room temperature, followed by permeabilization with 0.5% Triton X-100 for 15 min. Nuclei were stained with DAPI for 10 min and washed three times with PBS prior to imaging. For inducible U2OS cell lines, 4,000 cells/well were seeded and treated with doxycycline (1 µg/mL) for 48 or 72 h. For immortalized MEFs, 2,500 cells/well were seeded and treated with 4-hydroxytamoxifen (1 μM; Sigma, H7904) for 48 h. For rescue experiments, inducible U2OS lines and *Rpl11*^lox/lox^ MEFs were co-treated with rapamycin (50 nM and 100 nM, respectively) together with dox or 4-OHT as indicated. HTM images were acquired on an Opera Phenix™ Plus High-Content Imaging System (PerkinElmer) using a 10× objective. Cell numbers were obtained by automated DAPI-based nuclei segmentation and counting in Harmony (PerkinElmer).

### Microscopy

For HTM, cells were grown on μCLEAR-bottom 96-well plates (Greiner Bio-One). Cells were fixed in 4% paraformaldehyde (PFA) in PBS for 10 min at room temperature (RT), permeabilized with 0.5% Triton X-100 in PBS for 15 min at RT, and washed 3x with PBS. Samples were blocked for 30 min in PBS containing 2.5% bovine serum albumin (BSA). Primary antibodies were diluted in PBS/2,5% BSA and incubated or overnight at 4 °C, followed by 3x washes in PBS. Cells were incubated 1h at RT with secondary antibodies (Alexa Fluor–conjugated; Invitrogen/Thermo Fisher) diluted 1:5,000 in PBS and DAPI 0,5 µg/mL, followed by 3x PBS washes. Nascent RNA synthesis was monitored with 5-ethynyl uridine (EU, 10 µM, 45 min) and nascent protein synthesis with O-propargyl-puromycin (OPP, 20 µM, 45 min). Cells were then fixed and permeabilized as above, and the click-chemistry reaction was performed according to the manufacturer’s instructions (Click-iT Imaging Kits, Life Technologies/Invitrogen), followed by PBS washes and DAPI staining. Images were acquired automatically at 20x magnification on an Opera Phenix™ Plus High-Content Imaging System (PerkinElmer). Segmentation and quantification were performed in Harmony (PerkinElmer). DAPI-positive nuclei were used to generate nuclear masks. Mean fluorescence intensity of the indicated markers was extracted per cell (nucleus and/or cytoplasm as specified). Individual-cell measurements were exported and plotted; in the graphs, each dot represents a single cell. A full list of the reagents used in this study, including antibodies and the related information (supplier, catalogue number, dilution) is provided in **Table S2**.

### RNA FISH

Cells were grown on μCLEAR-bottom 96-well plates (Greiner Bio-One). Cells were fixed in 4% paraformaldehyde (PFA) in PBS (10 min) at room temperature (RT), permeabilized with 100% cold methanol (10 min) and rehydrated with 70% ethanol (10 min). Then, cells were incubated with 1M Tris pH8.0 for 5 min. The probe was prepared in hybridization buffer (1 mg/ml of Yeast tRNA, 0.005% BSA, 10% of dextran sulfate, 25% deionized formamide and 2X SSC) for a final concentration of 1 ng/ul and incubate overnight at 37 C maintained in humidity. After hybridization, cells were wash once with 4x SSC and once with 2x SSC, and then incubated with DAPI for 10 min. The sequence of the RNA probes (kindly provided by Angus Lamond) is detailed in **Table S2**.

### Western Blotting

Cell pellets were collected and washed with ice-cold PBS and lysed at 4 °C for 30 min in polysome lysis buffer (15 mM Tris-HCl pH 7.4, 15 mM MgCl_2_, 300 mM NaCl, 1% Triton-X-100, 0.1% β-mercaptoethanol, 200 U/mL RNAsin (Promega), 1 complete Mini Protease Inhibitor Tablet (Roche) per 10 mL). Lysates were clarified by centrifugation at 13,200 × rpm, 15 min, 4 °C, and protein concentration was determined with the Bio-Rad Protein Assay. Approximately, 30 µg of protein were mixed with NuPAGE LDS sample buffer (4x) and 10 mM DTT, heated 5–10 min at 95 °C, and loaded onto precast gradient gels (4–20% polyacrylamide; Thermo Fisher Scientific), and transferred using standard methods. After blocking, the membrane was incubated overnight at 4 °C with primary antibodies. The following day, membranes were incubated with a fluorophore-conjugated secondary antibody at RT, and protein detection was performed using the Li-Cor LCx system (Biosciences). For mTOR signaling analysis, U2OS cells were serum-starved overnight (O/N) in high-glucose DMEM supplemented with 0.2% FBS, 2 mM L-glutamine, and 1% penicillin/streptomycin. This starvation medium was maintained throughout the duration of the subsequent treatments until cell harvesting. Antibodies used are detailed in **Table S2**.

### Polysome analyses

Exponentially growing U2OS cells under the indicated treatments were incubated with 50 μg/mL of cycloheximide for ten minutes, washed with cold PBS and lysed in 1 mL of polysome buffer (Tris-HCl 30 mM pH 7.5, 100 mM KCl, 5 mM MgCl_2_, 1 mM DTT, 1% Triton X-100 and 50 μg/ml cycloheximide) for 10 min on ice. After three passages through a 22G needle, cell lysates were clarified by low-speed centrifugation and loaded on a 10-40% sucrose gradient prepared in polysome buffer. Gradients were centrifuged at 35,000 rpm in a SW-40 rotor for 3 h at 4 °C in a Beckman SW40.1 rotor and fractionated using an ISCO fractionator coupled to a UV recorder.

### Proteomics

Polysome-associated proteomics were performed as previously described^28^. In brief, samples were solubilized in 2% SDS, 100 mM TEAB pH 7.55. Proteins (20µg) were reduced and alkylated (15 mM TCEP, 25 mM CAA) 1h at 45 °C in the dark. Then, samples were digested following the solid-phase-enhanced sample-preparation (SP3) protocol^59^. Briefly, ethanol was added to the samples to a final concentration of 70% and proteins were incubated for 15 minutes with SP3 beads at a bead/protein ratio of 10:1 (wt/wt). Beads were rinsed using 80% EtOH and proteins were digested with 100 µl of trypsin in 50 mM TEAB pH 7.55 (promega, protein:enzyme ratio 1:50, 16 h at 37 °C). Resulting peptides were desalted using C18 stage-tips.

Liquid chromatography was performed using a Vanquish™ Neo HPLC system coupled to either an Orbitrap Astral (E01) or Orbital Exploris 480 (E03) (Thermo Fisher Scientific). Peptides were first loaded onto a trap column (PepMap™ Neo, 5 µm C18, 300 µm × 5 mm) and then separated on either an Aurora Elite TS (C18, 1.7 µm, 75 µm × 25 mm) (E01) or an Easy-Spray™ PepMap™ Neo analytical column (C18, 2 µm, 75 µm × 500 mm) (E03) both maintained at 50 °C. Peptides were separated at a flow rate of 400 nL/min, with Buffer A consisting of 0.1% formic acid (FA) and buffer B consisting of 100% acetonitrile (ACN) with 0.1% FA. The gradient ramped from 2% to 45% Buffer B over either 24 (E01) or 65 (E03) minutes, followed by a 5-minute wash at 98% Buffer B. Peptides were ionized using the EASY-Spray source at 1.8 kV (E01) or 1.5 kV (E03). The ion transfer tube was set to 300 °C.

MS1 were acquired at a resolution of 240,000. Precursor spectra were acquired every 0.6 seconds. Ion peptides were fragmented using higher-energy collisional dissociation (HCD) with a normalized collision energy of 27%. MS/MS spectra were acquired at a resolution of 80,000 (524 m/z). Normalized AGC target was set to 500%, with a maximum injection time of 3 ms for full MS and to 100%, with a maximum injection time of 3 ms for DIA MS/MS. Precursor isolation windows were set to 2m/z, with window placement optimized across 380-980 m/z. The precursor scan range was set to 390-980 m/z. Fragment ion scan range was set to 150-2000 m/z. The mass spectrometer was operated in a data-independent acquisition (DIA) mode using 60,000 precursor resolution and 15,000 fragment resolution. Ion peptides were fragmented using higher-energy collisional dissociation (HCD) with a normalized collision energy of 29. The normalized AGC target percentages were 300% for Full MS (maximum IT of 25 ms) and 1000% for DIA MS/MS (maximum IT of 22 msec). 8 m/z precursor isolation windows were used in a staggered-window pattern from 396.43 to 1004.70 m/z. A precursor spectrum was interspersed every 75 DIA spectra. The scan range of the precursor spectra was 390-1000mz.

For data analysis, raw files were processed with DIA-NN (1.8.2) using the library-free setting against a human protein database (UniProtKB/Swiss-Prot, 20,373 sequences) supplemented with contaminants. Afterwards, the “report.pg_matrix.tsv” file was loaded in Prostar^60^ using the intensity values for further statistical analysis. Briefly, proteins with less than three valid values in at least one experimental condition were filtered out. Missing values were imputed using the algorithms SLSA^61^ for partially observed values and DetQuantile for values missing on an entire condition. Differential analysis was done using the empirical Bayes statistics Limma. Proteins with a p.value < 0.01 and a log2 ratio >1 or <-1 were defined as regulated. The FDR was estimated to be below 2% by Benjamini-Hochberg.

### RNAseq

Total RNA was extracted using the Absolutely RNA Microprep kit (Agilent, 400805) following the manufacturer’s instructions. Libraries were prepared with the QuantSeq 3′ mRNA-Seq Library Prep Kit (Lexogen) and sequenced on an Illumina platform, yielding ∼10 million reads per sample. Raw sequence data in FASTQ format were processed using the *cluster_rnaseq* pipeline (https://github.com/cnio-bu/cluster_rnaseq). Initial quality control was performed with FastQC (v0.11.9) to assess read quality. Adapter sequences and low-quality bases were trimmed using BBDuk from the BBTools suite (v39.01). High-quality trimmed reads were aligned to the reference genome using STAR (v2.7.11b)^62^. For human samples, the GRCh38 primary assembly and GENCODE^63^ v43 annotation were used; for mouse samples, reads were mapped to the GRCm39 assembly with GENCODE vM37 annotation. Gene-level quantification was performed using featureCounts^64^ (v2.1.1) to generate a raw count matrix. Quality control metrics from all processing stages were aggregated and visualized using MultiQC^65^ (v1.18). Ortholog mapping and gene ID conversions across human and animal models were performed using the biomaRt R package^66^, utilizing the Ensembl database as the primary reference for cross-species annotations. Lowly expressed genes were removed using the filterByExpr function from the edgeR R package, which applies a minimum count threshold based on library sizes and experimental group sizes.

### Covariate selection and statistical adjustment of transcriptomic data

To prevent confounding of biological signals, we implemented a structured covariate selection framework for each dataset, applied both to differential expression (DE) regression models and to log-CPM matrix correction via limma’s^67^ removeBatchEffect for visualization purposes. For the NYGC ALS Consortium and blood sample datasets, collinearity among metadata variables was first assessed using Cramér’s V (categorical) and Spearman’s correlation (numerical), excluding redundant predictors (threshold > 0.7). Following TMM normalization and log2-CPM transformation, PCA was performed and the top ten PCs were regressed against candidate covariates to estimate variance explained (R²). variancePartition^68^ was used as a complementary approach to quantify per-variable contributions to total transcriptional variance. Final covariates were selected based on variance explained or established biological relevance in ALS; the latter criterion ensured inclusion of Age and Sex regardless of their statistical prominence in specific cohorts. Covariates per dataset were as follows: for the NYGC ALS Consortium (all tissues), library preparation method, Sex, Age at death, and Age² (squared term for Age at death was incorporated to account for potentially non-linear dependencies); for blood samples, GC content, RNA concentration, RIN, Sex, Age at collection, and Age². For the GTEx dataset^69^, the high dimensionality of technical confounders prompted the use of Surrogate Variable Analysis (SVA)^70^, following a previously established strategy for this resource^71^. The number of surrogate variables was estimated with num.sv() using the ‘be’ method^72^, with Age and Sex protected in the full model to ensure SVs captured only technical variation. Due to the controlled laboratory environment and the absence of significant batch effects in the PCA, no additional covariates were incorporated for the remaining datasets.

### Differential gene expression analyses

DE analysis was conducted using the edgeR R package^73,74^. To maximize statistical power and reduce noise, lowly expressed genes were first filtered out using the filterByExpr function, which determines a filtering threshold based on the experimental design and library sizes. Normalization factors were then calculated using the TMM method to account for compositional biases between libraries. Statistical testing was performed within the Quasi-likelihood framework. For each dataset, the design matrix incorporated the biological variables of interest alongside the specific technical and clinical covariates previously identified. Differentially expressed genes were selected using a significance threshold of p-value < 0.05.

### Enrichment analyses

To evaluate pathway activity at the single-sample level, enrichment scores were generated using the GSVA R package^75^ implementing the ssGSEA method. These scores were calculated from TMM-normalized log2-CPM matrices, which were adjusted for the previously identified technical artifacts and clinical covariates via limma’s removeBatchEffect function. Pathway gene sets used in these analyses were retrieved from The Molecular Signatures Database (MSigDB)^76^ and the Jensen curated collection^77^, or manually established based on relevant literature to target specific disease signatures. The resulting enrichment scores were subsequently used for visualization and statistical comparisons between conditions, where significant differences in pathway activity were evaluated using Student’s t-tests.

To identify changes in biological pathways, Gene Set Enrichment Analysis (GSEA)^78^ was performed using the fgsea R package^79^. Genes were ranked by their log2FC values from the differential expression analysis. Pathway collections were retrieved from MSigDB via the msigdbr package, including Hallmark (H), Curated (C2) and Gene Ontology (C5), as well as custom curated signatures from Jensen. Statistical significance was determined by calculating Normalized Enrichment Scores (NES) to aid comparison between sets.

### Definition and analysis of the ALS^sig^

An initial ALS candidate signature was established by intersecting significantly up-regulated genes (p-value < 0.01, logFC > 0.5) across the three transcriptomic datasets derived from U2OS cells expressing (PR)_97_, hnRPNA2^D290V^ or TDP43^A315T^. The NYGC Cervical Spinal Cord dataset served as the discovery cohort for signature refinement. The resulting normalized and corrected expression matrix for the aforementioned technical and clinical covariates was used for subsequent modeling and feature selection. To identify the most informative genes within the initial candidate list, we applied the Boruta algorithm^80^, a feature selection wrapper based on Random Forest^81^ that evaluates variable relevance against randomized “shadow” features. To address class imbalance, the underlying Random Forest model (1,000 trees, 500 runs) employed stratified downsampling to ensure balanced representation of ALS and Control samples across trees. Genes confirmed as “Important” by Boruta constituted the Refined ALS Signature, while the remaining independent datasets were reserved as validation cohorts to assess cross-tissue robustness.

The generalizability of the Refined ALS Signature was assessed across all available NYGC ALS Consortium tissues (Spinal Cord Cervical, Spinal Cord Lumbar, Cerebellum, Frontal Cortex, Lateral Motor Cortex, and Medial Motor Cortex) using ROC curve analysis (pROC R package)^82^ on ssGSEA enrichment scores. AUC was calculated per tissue independently; Spinal Cord Cervical served as the discovery cohort, with all remaining tissues as independent validation sets. For benchmarking, the same analysis was applied to the KEGG ALS pathway and Jensen ALS gene sets.

The association between the ALS gene signature and prognosis was assessed in ALS patients from the cervical spinal cord cohort with available disease duration data. Disease duration (months from symptom onset to death) served as the survival endpoint; as all samples were post-mortem, no censoring was applied (vital status = 1). Patients were stratified into high and low ssGSEA score groups using percentile-based cutoffs, with intermediate cases excluded to maximize group contrast. Kaplan-Meier survival curves were estimated using the *survival* R package^83^ and visualized with *survminer*^84^ (v0.4.9); group differences were assessed by log-rank test.

### Data availability

Mass spectrometry data have been deposited to the ProteomeXchange Consortium via the PRIDE partner repository with the dataset identifier PXD071379. RNA-seq data are available at the GEO repository with accession number GSE327905. External RNAseq datasets used in this study include: NYGC ALS Consortium (GSE153960); bulk RNAseq of ALS spinal cords (GSE287256) or blood samples (GSE234297) from independent datasets; DBA lymphoblastoids (GSE119954).

### Statistics

All statistical analyses were performed using Prism software (GraphPad Software) and statistical significance was determined where the p-value was <0.05 (*), <0.01 (**), <0.001 (***) and <0.0001 (****).

## Acknowledgments

The authors want to thank Manuel Serrano for sharing the *Rpl11*^lox^ mouse strain, Angus Lamond for providing the sequence of the probes used for rRNA FISH, Daniela Hühn for comments on the manuscript and the proteomics, genomics, confocal microscopy and transgenics units of the CNIO for their technical support in this study. Work done in the O.F-C. lab was supported by grants from the Swedish Research Council (VR) (538-2014-31), the Hjärnfonden foundation (FO2023-0263) the Spanish Ministry of Science, Innovation and Universities (PID2024-163103OB-I00, co-financed with European FEDER funds), La Caixa Foundation (HR22-00890) and CIBERNED (CB24/05/00022).

## Author Contributions

A.S. contributed to most of the experiments and to the preparation of the figures; G.V. contributed to bioinformatic analyses; A.E.-M. helped with some of the experiments on ribosomal proteins and ribosomopathy models; I.V. helped with polysome fractionations; S.R. provided technical help; V.L. performed experiments and contributed to the experimental plan and supervision of the study; O.F. conceived the experimental plan, coordinated the study, supervised the experiments and wrote the manuscript.

