## Supplementary material for "The accumulation of orphan ribosomal proteins is a hallmark of ALS": Figures S1-2; Tables S1-2

Saez-Mas et al **Fig. S1**

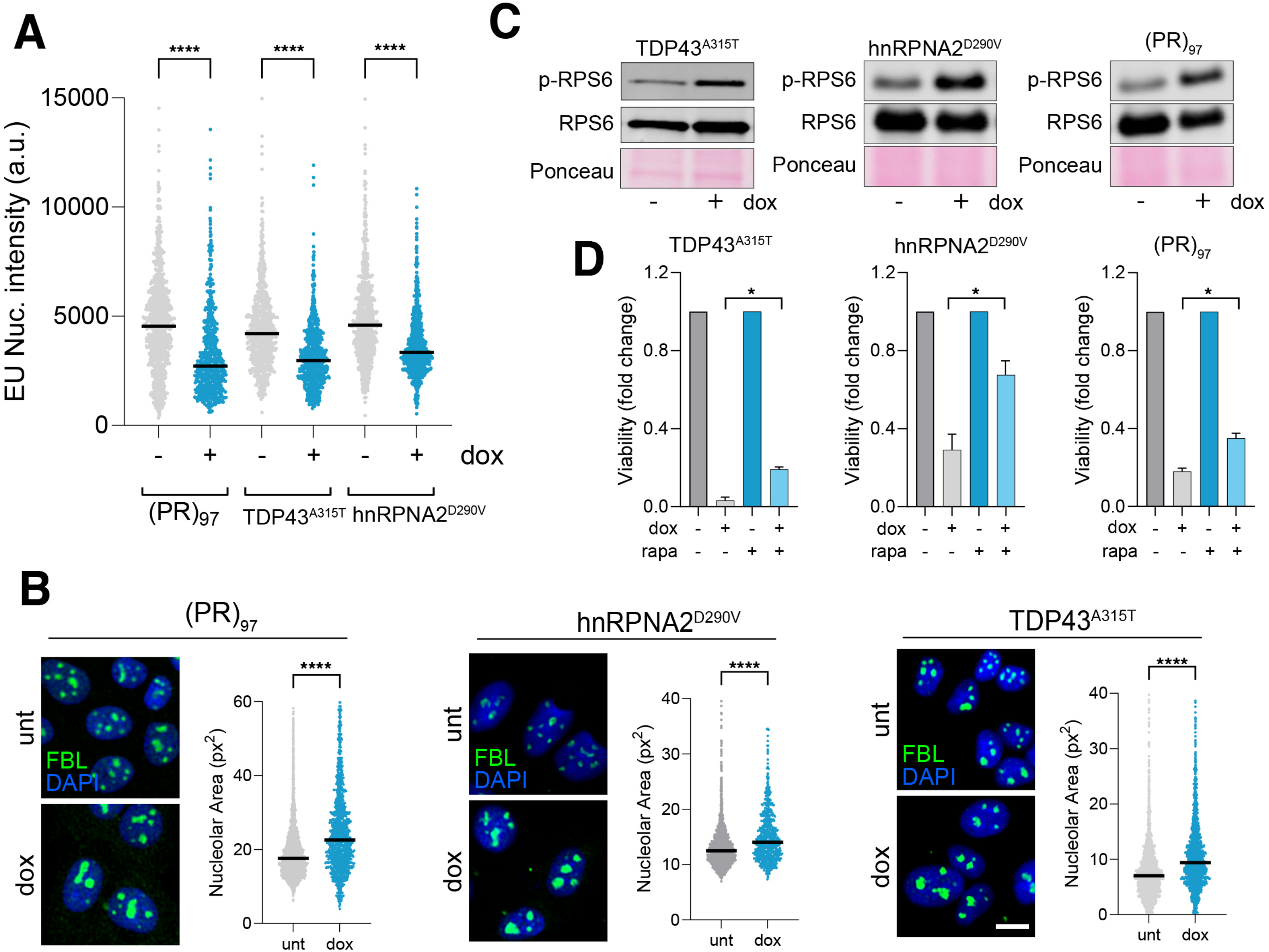

**Figure S1. Nucleolar and mTOR dynamics upon expression of ALS mutations.** (**A**) HTM-dependent quantification of RNA synthesis (as monitored by the incorporation of 5´-Ethynyluridine (EU)), in U2OS cells expressing $(PR)_{97}$, $hnRPNA2^{D290V}$ or $TDP43^{A315T}$. Cells were treated with dox for 48h, followed by a 1h pulse with EU. Black lines indicate median values. The graph represents one experiment from 3 biological repeats. ****$P < 0.0001$; *t*-test. (**B**) HTM-dependent quantification of the nucleolar area (as monitored by the FBL signal (green)), in U2OS cells expressing $(PR)_{97}$, $hnRPNA2^{D290V}$ or $TDP43^{A315T}$. Black lines indicate median values. DAPI was used to stain nuclei. Example images are provided to the left of the corresponding quantification. Scale bar (white) represents 5 μm. The graph represents one experiment from 3 biological repeats. ****$P < 0.0001$; *t*-test. (**C**) WB to analyse RPS6 phosphorylation (p-RPS6) levels in U2OS cells expressing $(PR)_{97}$, $hnRPNA2^{D290V}$ or $TDP43^{A315T}$. Total levels of RPS6 are shown as a loading control, together with the Ponceau image from the WB. (**D**) Cell viability, calculated by HTM-dependent quantification of nuclear numbers, in U2OS cells expressing $(PR)_{97}$, $hnRPNA2^{D290V}$ or $TDP43^{A315T}$, in the presence or absence of rapamycin (rapa: 50 nM). The graph represents one experiment from 3 biological repeats. *$P < 0.05$; *t*-test.

**A**

| Tissue Region | Samples | | | ALS mutations | | | | |
|---|---|---|---|---|---|---|---|---|
| ***Tissue Region*** | ***Total*** | ***Control*** | ***ALS*** | ***Unknown*** | ***C9ORF72*** | ***SOD1*** | ***OPTN*** | ***Others*** |
| *Spinal Cord (Cervical)* | 235 | 46 | 189 | 149 | 31 | 3 | 1 | 5 |
| *Spinal Cord (Lumbar)* | 219 | 41 | 178 | 142 | 26 | 3 | 2 | 5 |
| *Cerebellum* | 207 | 52 | 155 | 123 | 24 | 4 | 1 | 3 |
| *Frontal Cortex* | 255 | 67 | 188 | 146 | 32 | 3 | 2 | 5 |
| *Lateral Motor Cortex* | 133 | 27 | 106 | 84 | 17 | 3 | 0 | 2 |
| *Medial Motor Cortex* | 133 | 30 | 103 | 81 | 15 | 4 | 1 | 2 |
| ***Total number*** | **1182** | **263** | **919** | **725** | **145** | **20** | **7** | **22** |

**B**

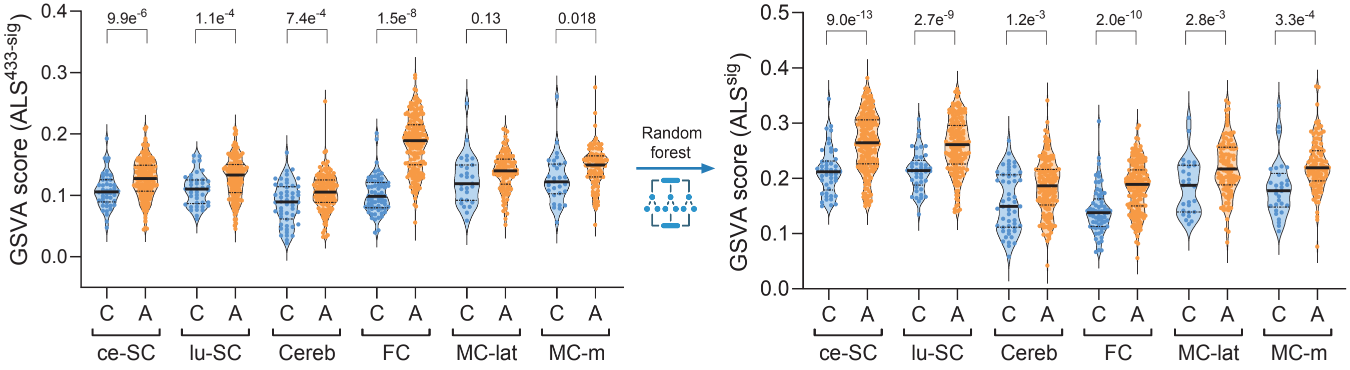


**C**

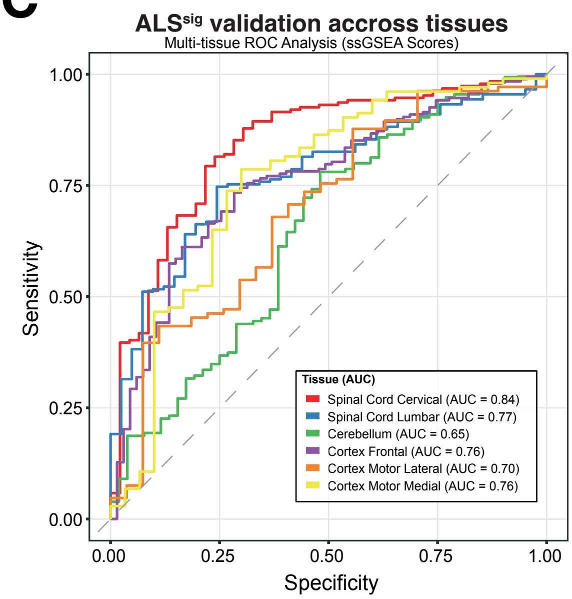


**D**

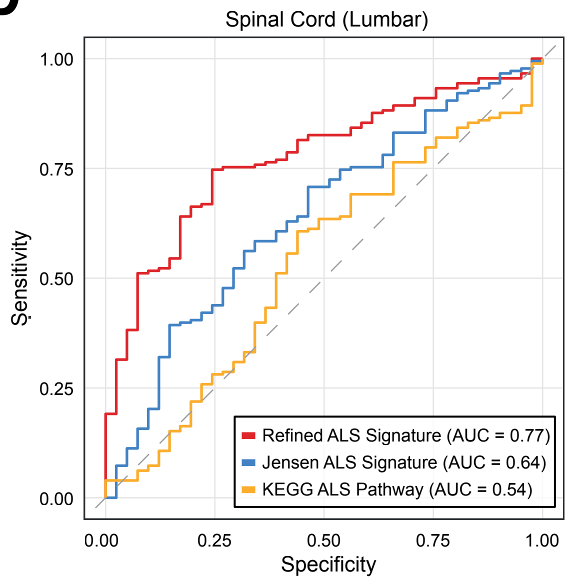

**Figure S2. Validation of the ALS$^{sig}$ across different tissues and datasets.** (**A**) Summary of the NYGC ALS cohort metadata, including the number of samples per tissue, distribution between control and ALS cases, and the number of samples harboring the specified mutations. (**B**) Gene Set Variation Analysis (GSVA) scores for the initial ALS signature (based on 433 genes, prior to machine learning refinement) and the final ALS$^{sig}$ across different tissues of the NYGC dataset. (**C**) ROC curve showing the diagnostic performance (AUC) of the ALS$^{sig}$ across different tissues of the NYGC dataset. (**D**) ROC curve comparing the diagnostic performance (AUC) of the refined ALS$^{sig}$ with established ALS-related signatures from the Jensen and KEGG databases in lumbar spinal cord simples.

**TABLE S1.** GENES FROM THE ALS$^{SIG}$.

LDAF1
FAS
SLAMF7
ASB1
FAM107B
MAST4
SLC6A15
DAAM1
MOSPD1
PBDC1
RIPK2
PANX1
CLEC2B
ELK3
PHACTR1
GNPDA1
SSR3
PRRX1
ARHGEF2
DNAJC6
RBM18
FXR2
DPH2
DCLK1
SPX
RSL24D1
HECW2
AP1AR
SINHCAF
DUSP10
PPP1R18
PLIN2
GSTO1
ENKUR
CCDC148
CEBPG
LY96
C21orf91
ANTXR2
MON1A
STEAP1
HEY1
MMADHC
TMC7
GNG12
NIPAL4
GLRX
TRIB1
NABP1
ASPHD1
SH3BP5L
DLGAP1-AS1
AFMID
ZNF267
PPTC7
CASP4
VEPH1
PDCD1LG2
HLA-E
PTCHD4
TXNIP

**TABLE S2.** LIST OF REAGENTS USED IN THIS STUDY.

| REAGENT or RESOURCE | SOURCE | IDENTIFIER |
|---|---|---|
| **Antibodies** | | |
| anti-FBL | CST | 2639 |
| anti-RPL11 | Proteintech | 14583-1-AP |
| anti-HA | Roche | 11867423001 |
| Anti-RPS6 | CST | 2217 |
| anti-p-RPS6 (Ser235/236) | CST | 2215 |
| anti-ACTIN | Sigma | A5441 |
| anti-TUBULIN | Sigma | T9026 |
| anti-VINCULIN | Abcam | ab129002 |
| anti-Mouse-Alexa647 | Thermo | A21463 |
| anti-Rabbit-Alexa488 | Thermo | A21441 |
| anti-Rabbit-Alexa555 | Bethyl | A120-201D3 |
| **Chemicals, peptides, and recombinant proteins** | | |
| Blasticidin | Life Technologies | A11139-03 |
| Zeocin | Gibco | R25005 |
| Doxycycline | Sigma | D9891 |
| 4-hydroxytamoxifen | Sigma | H7904 |
| Rapamycin | Alpha Aesar | J62473 |
| Tetracycline-free FBS | PAN-Biotech | P30-3602 |
| Lipofectamine 2000 | Thermo Fisher | 11668027 |
| DAPI | Invitrogen | D1306 |
| **Commercial kits** | | |
| Click-iT OPP Alexa 647 Fluor Protein Synthesis Assay | Thermo Scientific | C10458 |
| Click-iT RNA Alexa 488 Fluore Alexa Fluor Assay | Thermo Scientific | C10329 |
| RNA Microprep kit | Agilent | 400805) |
| QuantSeq 3' mRNA-Seq Library Prep Kit | Lexogen | N/A |
| **Deposited data** | | |
| Polysome proteomics data | This study | PRIDE: PXD071379 |
| RNA-Seq data | This study | GEO: GSE327905 |
| **Experimental model: Cell lines** | | |
| *Rpl11*$^{+/+}$ and *Rpl11*$^{lox/lox}$ MEF | Morgado-Palacin et al. 2015 | |
| U2OS | ATCC | HTB-96 |
| U2OS- HNRNPA2$^{D290V}$ (TetON) | This study | |

**TABLE S2.** LIST OF REAGENTS USED IN THIS STUDY.

| | | |
|---|---|---|
| U2OS- TDP43$^{A315T}$ (TetON) | Colicchia et al. 2022 | |
| U2OS-PR97 (TetON) | Sirozh et al. 2024 | |
| **Probes** | | |
| 47S pre-rRNA: 5´-AGA CGA GAA CGC CTG ACA CGC ACG GCA C -3´– CY3 | Kind gift from A. Lamond | |
| 28S rRNA : 5´- GAG GGA ACC AGC TAC TAG ATG GTT CGA TTA-CY3. | Kind gift from A. Lamond | |
| **Recombinant DNA** | | |
| pJ4M_hnRNPA2_FL_D290V | Addgene | 139110 |
| pINTO-HNRNPA2$^{D290V}$-HA-flag | This study | |
| **Software and algorithms** | | |
| Harmony | | https://www.revvity.com |
| GraphPad Prism | Version 11 | https://www.graphpad.com |
| R Studio | Version 4.4.0 | https://rstudio-desktop.softonic.com/ |
| Gene Set Enrichment Analysis (GSEA) | Version 4.4.0 | https://www.gsea-msigdb.org/ |
| Cluster_RNAseq | | https://github.com/cnio-bu/cluster_rnaseq |
| STRING | | https://www.string-db.org |
